# Analyses of cancer incidence and other morbidities in neutron irradiated B6CF1 mice

**DOI:** 10.1101/2020.03.26.009852

**Authors:** Alia Zander, Tatjana Paunesku, Gayle Woloschak

## Abstract

The Department of Energy conduced ten large-scale neutron irradiation experiments at Argonne National Laboratory between 1972 and 1989. Using a new approach to utilize experimental controls to determine whether a cross comparison between experiments was appropriate, we amalgamated data on neutron exposures to discover that fractionation significantly improved overall survival. A more detailed investigation showed that fractionation only had a significant impact on the death hazard for animals that died from solid tumors, but did not significantly impact any other causes of death. Additionally, we compared the effects of sex, age first irradiated, and radiation fractionation on neutron irradiated mice versus cobalt 60 gamma irradiated mice and found that solid tumors were the most common cause of death in neutron irradiated mice, while lymphomas were the dominant cause of death in gamma irradiated mice. Most animals in this study were irradiated before 150 days of age but a subset of mice was first exposed to gamma or neutron irradiation over 500 days of age. Advanced age played a significant role in decreasing the death hazard for neutron irradiated mice, but not for gamma irradiated mice. Mice that were 500 days old before their first exposures to neutrons began dying later than both sham irradiated or gamma irradiated mice.

## INTRODUCTION

Ionizing radiation can be classified by its linear energy transfer (LET) to better understand how quickly radiation is attenuated and the concentration of energy deposited near the particle track. Experiments evaluating the biological effects of low and high LET ionizing radiation found that high LET radiation is more damaging to biological material in part because sites of DNA double strand damage and other types of damage are in closer proximity to one another than occurs with low LET radiation. These clusters of damage make DNA repair more difficult and lead to increased cell death (1). Neutrons are the smallest and the most penetrating of all particles included in the category of high LET ionizing radiation. Neutrons are further classified by the amount of kinetic energy in a free neutron and this energy ranges from less than 0.02eV for cold neutrons to over 20MeV for ultrafast neutrons. While numerous animal studies were conducted in the USA, Europe and Asia using low LET radiation such as x-rays and gamma rays (2–5), extensive neutron irradiation experiments were less common.

Between 1972 and 1989, Argonne National Laboratory (ANL) housed one of the few neutron irradiators called the JANUS reactor suitable for large-scale whole-body animal irradiations with fission spectrum neutrons with an energy range peak at 1 MeV (6–10). During this time, ANL performed ten independent experiments investigating the effects of gamma and neutron irradiations on mice. (11). These animals were irradiated under different conditions and allowed to live out their entire lifespan. Ionizing radiation exposures ranged from low to high doses, included fractionated and acute exposures, and the age when mice were first exposed to ionizing radiation varied by experiment. Moribund mice were necropsied to determine a cause of death (COD), other death contributing diseases, and non-contributing diseases. The resulting dataset from the Janus studies includes information on over 50,000 male and female mice (3, 11–13). A neutron irradiation study of this magnitude will most likely never be reproduced and it is important that researchers continue to analyze these data in light of novel biological findings and analytical methods. Innovative statistical approaches, such as the ones used here, can lead to new insights into the effects of neutron irradiation on whole organisms.

In this study we analyzed neutron irradiated mice using methods similar to those used for investigation of cobalt 60 gamma irradiated mice [Zander et al. submitted]. Here, we included comparisons between key findings from our analyses on neutron and gamma irradiated mice. Comparisons between the two qualities of radiation are beneficial for determining the underlying biological factors leading to differential health outcomes. Numerous *in vivo* studies comparing x-rays or gamma rays with neutrons evaluated life shortening or focused on genetic changes to establish a suitable relative biological effectiveness factor (RBE) (6–9, 14–20). Work with radioprotectors and neutron irradiation *in vivo* and *in vitro* also found profound differences between the high and low LET ionizing radiation (21–24).

Previous neutron irradiation studies suggested that fractionation had little effect on overall survival or cancer risk from neutron exposures (6, 7, 9, 25–30). We re-analyzed data from the Janus experiments conducted at ANL by pooling results from different studies in a new way in order to determine the role of fractionation in neutron irradiations for overall survival and for risks of specific causes of death. Rigorous statistical testing on sham irradiated mice enabled us to combine several Janus experiments into one large dataset. This approach increased our statistical power and allowed comparisons of different fractionation regimens across a wide array of total doses. [Zander et al., submitted]. A more voluminous dataset enabled by this approach allowed us to explore neutron fractionation effects in young and old mice across a wide array of total doses of fission spectrum neutrons.

## METHODS

### Data Selection

Details on data selection methods used in this study can be found in our previously published work [Zander et al., co-submitted]. To increase the statistical power given the limited set of fractionation schedules for neutron irradiated mice, we combined mice that received their total doses in 24 and 60 fractions into a single group. Therefore, in addition to the initial analysis of control mice, we also verified that grouping together 24 and 60 sham irradiation fractions did not significantly change survival probabilities in control mice compared to animals that received acute sham exposures. Because the greatest number of fractions a neutron irradiated mouse received was only 60 fractions, we also excluded all animals with more than 60 sham irradiation fractions from this work. We used Cox proportional hazards models with sex and a categorical fractionation term as independent variables for the main model **(S1 Fig A and B)**. For sensitivity analysis, we also stratified by sex **(S 1 Fig C and D)** and found no significant changes in the model output. Finally, we used Kaplan Meier survival curves to validate the proportional hazards assumption in our model **(S1 Fig E)**. After filtering out any control mice that received more than 60 sham irradiation fractions, we again verified that there were no significant differences between control mice based on experiment number or age first irradiated. **S1 Table** outlines the filtering process used to determine an acceptable cohort of neutron irradiated mice that combined data from multiple Janus experiments.

### Survival Analysis

Kaplan-Meier (KM) curves were utilized for categorical univariate survival analysis in R through the survfit function within the survival package (31, 32). Cox proportional hazard (PH) models were utilized to evaluate survival over time for multivariate models. Cox PH models enabled us to include interactions between variables, incorporate quantitative and categorical variables, and stratify by nuanced variables (33). The main Cox PH model used for neutron irradiated mice was as follows:

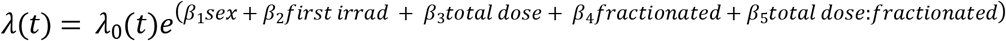

where *λ*(*t*) is the hazard function based on our set of covariates (sex, age first irradiated, total dose, fractionated (acute vs. fractionated), and the interaction between total dose and fractionated), ***β*** is a vector of their associated parameter estimates, and *λ*_0_(*t*) is the baseline hazard. All Cox PH models were created in R through the coxph function from the survival package (32).

### Competing Risks Analysis

We conducted competing risks analyses to understand the risks and probabilities of specific causes of death. In situations where there are no competing risks, or events that could decrease the likelihood of observing the event of interest (i.e. other causes of death), the cumulative incidence of an event is calculated as the one minus the survival function estimated with KM curves. However, when competing risks are present, the KM method is biased upward and the Fine and Gray method for CIF is more appropriate for examining probabilities of particular event types (34). For our competing risks analyses, we assessed crude incidences, cause-specific hazard models, and cumulative incidence function (CIF) regression models (also known as the subdistribution hazard) (35). To investigate crude, nonparametric incidences with competing risks present, we utilized the cuminc function from the cmprsk package in R (36)

We utilized both cause specific hazards and CIF models for multivariate regression analyses in the presence of competing risks. Cause specific hazards are used to determine the effect that covariates have on all event free subjects. These hazards were estimated using the coxph function from the survival package in R (32). For this method all causes of death other than the event of interest, were censored. Concretely:

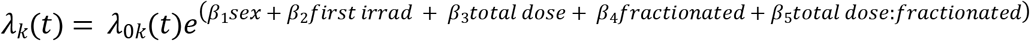,subset data for age first irradiated < 500 days.

where *λ*_*k*_(*t*) is the hazard function for the k^th^ cause of death.

The cumulative incidence function describes the overall likelihood of a specific result and does not depend on a subject being event free (34, 37–39). We utilized the crr function in the cmprsk package in R to model CIF regression hazards (36). Concretely:

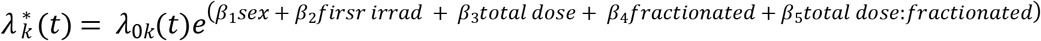,subset data for age first irradiated < 500 days.

where 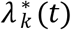 is the subdistribution hazard function for the k^th^ cause of death.

### Cause of Death Animal Groupings

The full list of causes of death separated by radiation treatment conditions (control, gamma rays, neutrons) and subdivided by gender is shown in **S2 Table**. “Grouped Macros” data was downloaded from the Janus website and includes all pathologies found at necropsy. CODs were categorized as lethal (L), contributory (C), or non-contributory (N) (11). For the purposes of this study, we only investigated lethal disorders. We grouped causes of death as solid tumors other than lymphomas (referred to as tumors), lymphomas, non-tumors, or cause of death unknown (CDU) to gain a stronger understanding of disease trends. For certain analyses we also looked at lung tumors separately from all other non-lymphoma solid tumors and examined non-thymic lymphoma specifically.

### Survival Analysis for JM8 mice

The main Cox PH model used for gamma irradiated JM8 mice analysis was:

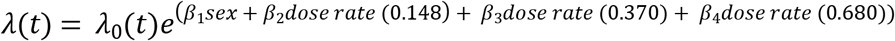, all mice included.

The main Cox PH model used for neutron irradiated JM8 mice analysis was:

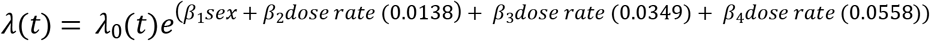, all mice included.

## RESULTS

### Fractionation decreases the death hazard in mice exposed to neutrons

We examined age at death in mice from Janus experiments with different total doses and fractionation regimens (**Fig 1**). The range of total doses for neutron irradiated mice was 0.94cGy - 301cGy. A minor subset of animals was first exposed to neutrons when they were over 500 days old, while the remainder of the mice were first irradiated around 100 days +/− 15 days **(Fig 1B)**. The highest acute exposure dose was 37.68cGy for young mice and 150.72cGy for aged mice. When comparing the age at death for aged mice compared to mice first irradiated around 100 days, there was a noticeable increase in longevity in the aged irradiated mice. No equivalent finding was present in the gamma irradiated young and old mice cohorts **(Fig 1C and Zander, submitted Fig 2C)**. We excluded the aged irradiated mice from our main analysis because none of the animals in our cohort were aged before sham irradiations and this small subset of data would have had a large amount of leverage on our model. Aged mice were, however, incorporated in robustness tests. Excluding these 560 aged mice from further investigation resulted in 14,018 mice total for our neutron analysis.

**Fig 1:**
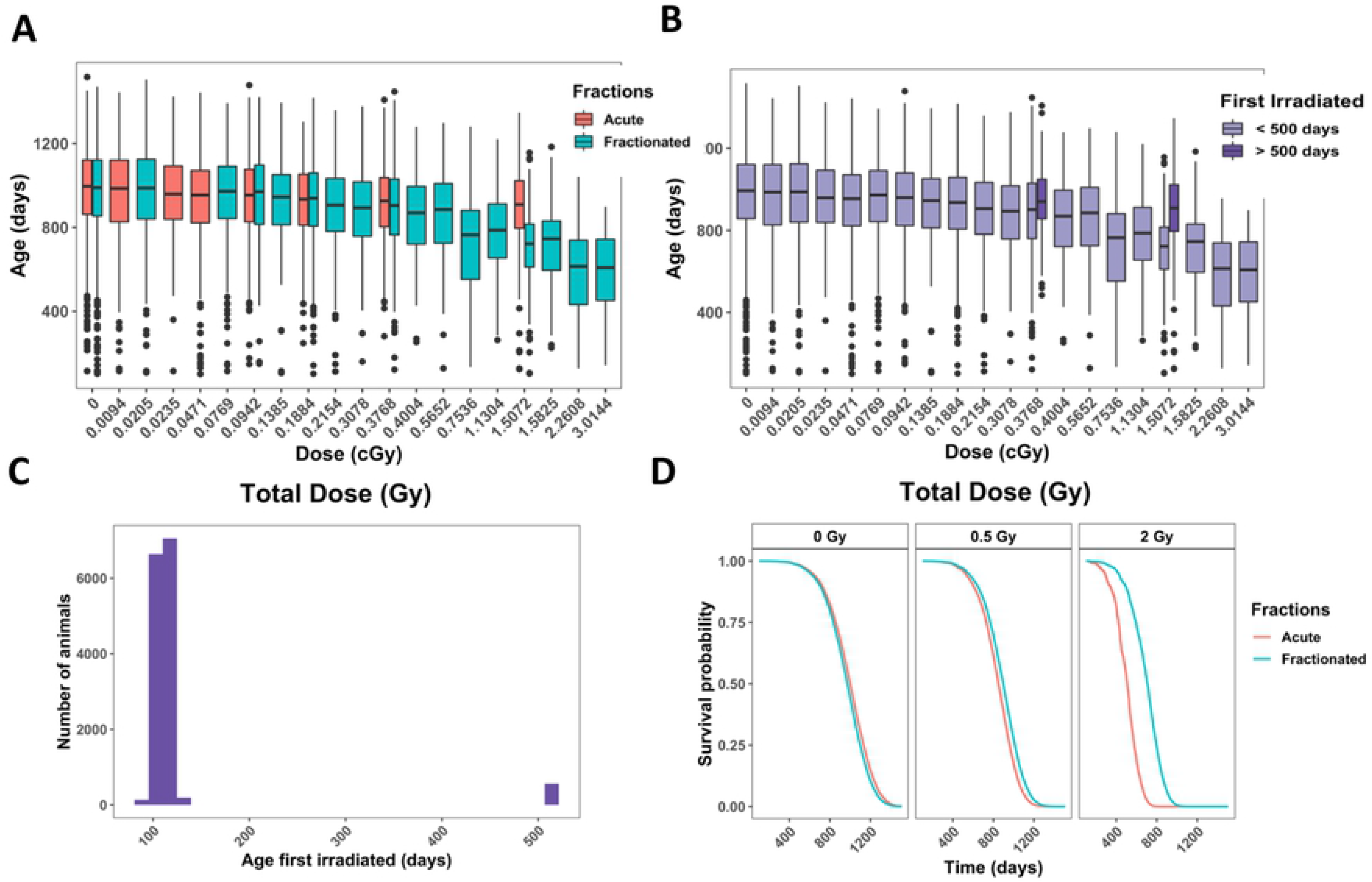
Analysis on animals filtered in Table 1 that were controls or neutron irradiated. **(A)** Box plot of age at death in days versus total dose in Gy. Colors indicate the number of fractions. **(B)** Histogram of the total number of animals versus age first irradiated in days. **(C)** Age at death in days versus total dose in Gy. Colors indicate whether a mouse was first irradiated before or after 500 days. **(D)** Representative graphs from Cox PH model output with age at death as the time scale and sex, age first irradiated, total dose, fractions, and the interaction between total dose and fractions as independent variables. The predicted outcomes shown are for female mice first irradiated at 120 days. All independent variables were significant in the model, as shown by the parameter estimates and statistical output from the model **(E)**.

**Table 1:**
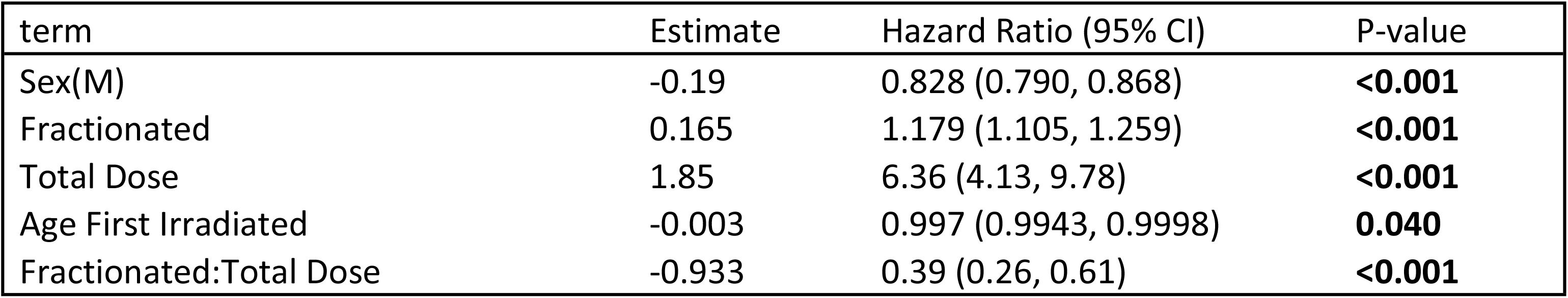
Parameter estimates, hazard ratios with 95% confidence interval, and p-values for main Cox

| term | Estimate | Hazard Ratio (95% CI) | P-value |
| --- | --- | --- | --- |
| Sex(M) | -0.19 | 0.828 (0.790, 0.868) | **<0.001** |
| Fractionated | 0.165 | 1.179 (1.105, 1.259) | **<0.001** |
| Total Dose | 1.85 | 6.36 (4.13, 9.78) | **<0.001** |
| Age First Irradiated | -0.003 | 0.997 (0.9943, 0.9998) | **0.040** |
| Fractionated:Total Dose | -0.933 | 0.39 (0.26, 0.61) | **<0.001** |

The relative biological effectiveness (RBE) between gamma rays and neutrons based on life shortening per cumulative dose was estimated to be around 10 for doses up to 40cGy and approximately 5 or less for neutron doses over 40cGy [(11)– page 36]. Because of this RBE estimate, the maximal total dose for neutrons was approximately one order of magnitude lower than the maximum dose used for gamma irradiations. In addition, the fractionation schedule was more limited and neutron irradiated mice only received 24 or 60 fractions. We combined these two fractionation conditions to increase sample size.

Using Cox PH models to study the effects of fractionation on survival in neutron irradiated mice, we found that the key interaction term between fractionations and total dose significantly decreased the death hazard **(Fig 1D and Table 1)**. Similar to gamma irradiated mice, males had a significantly lower death hazard compared to female mice and the main effects for fractionation and total dose significantly increased the hazard. Even with a limited range of ages from 90 to 180 days, an increase in age first irradiated significantly decreased the hazard. All of these variables showed statistical significance in our model, including the age first irradiated, a variable that was not significant in gamma irradiated mice **[Zander et al., submitted,]**. Robustness tests showed similar results, increasing our confidence in the benefit of fractionation on neutron irradiated mice **(S3 Fig)**. When we included aged mice for our robustness testing, acute exposures appeared to be less hazardous compared to fractionated exposures **(S2 Fig A and 2B)**. However, after correcting the model by adding in an interaction term between age first irradiated and total dose to account for the increased rescue effect of older ages at higher doses **(Fig 1C)**, fractionation decreased the death hazard **(S2 Fig C and 2D)**. Adding an interaction term between age first irradiated and total dose for the main model did not change the significance nor the sign of any terms in the model, and the interaction term itself was not significant **(S2 Fig E and 2F)**. Stratifying by sex **(S2 Fig G and H)** and including exact fractionation regimen and age first irradiated for control mice **(S2 I and J)** had no major consequence on the model. We used KM curves to validate the proportional hazards assumption in our model and found parallel survival curves between groups based on sex, number of fractions, age first irradiated, and total dose **(S3 Fig)**.

### In neutron irradiated mice, fractionation significantly decreases the hazard for tumor deaths

Accounting for competing risks by calculating cause-specific hazards, we found that the interaction between total dose and the fractionation variable was only significant for tumor deaths **(Fig 2A and Table 2)**. The interaction was beneficial for survival in groups of animals that developed tumors and lymphoma **(Fig 2B and Table 2)** and detrimental for mice dying of non-tumors and CDU **(Fig 2C and 2D and Table 2)**. However, this difference was only statistically significant for tumors. To evaluate this further, we inspected death due to lung tumors **(S4 Fig A)**, non-thymic lymphoma **(S4 Fig C)**, and tumors other than lung tumors **(S4 Fig B)**. The interaction term between fractionation and total dose was not significant for lung tumors nor non-thymic lymphoma, but was still significant in all tumors excluding lung tumors **(S4 Fig and Table 2)**. Males had a lower death for all individual causes of death, except for tumors. Males had a higher hazard for lung tumors specifically, but not for tumors excluding lung tumors, indicating that lung tumor risk in males was driving the entire outcome for the tumor group.

**Fig 2:**
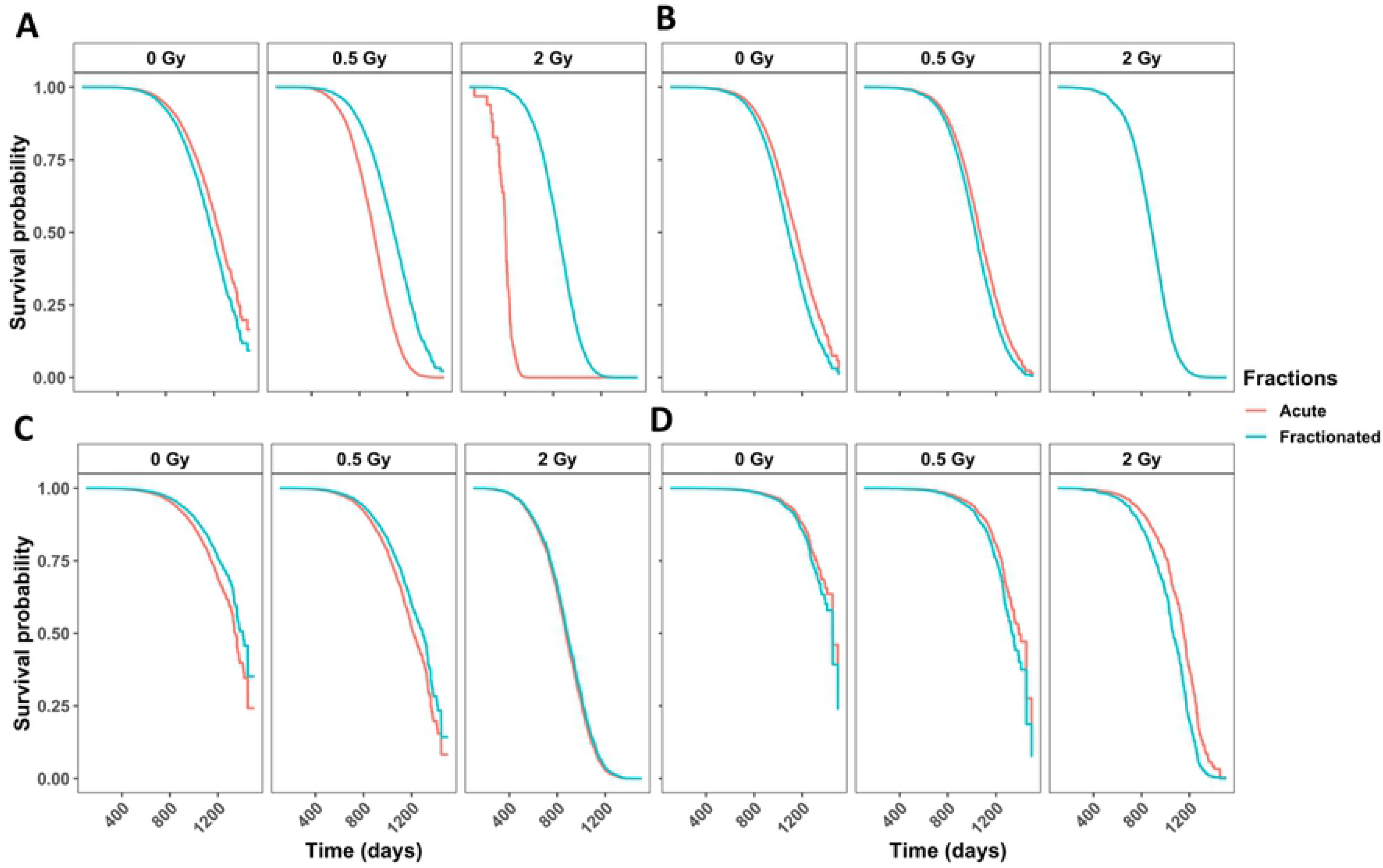
Competing risks models for specific causes of death in neutron-irradiated mice with age as a time scale and sex, age first irradiated, total dose, fractions, and the interaction between total dose and fractions as independent variables. Survival curves for cause of death being **(A)** any solid tumors, **(B)** lymphoma, **(C)** non-tumors, and **(D)** cause of death unknown. Model estimates, confidence intervals and p-values are listed in **Table 2**. The graphs represent predicted outcomes for female mice first irradiated at 120 days.

**Table 2:**
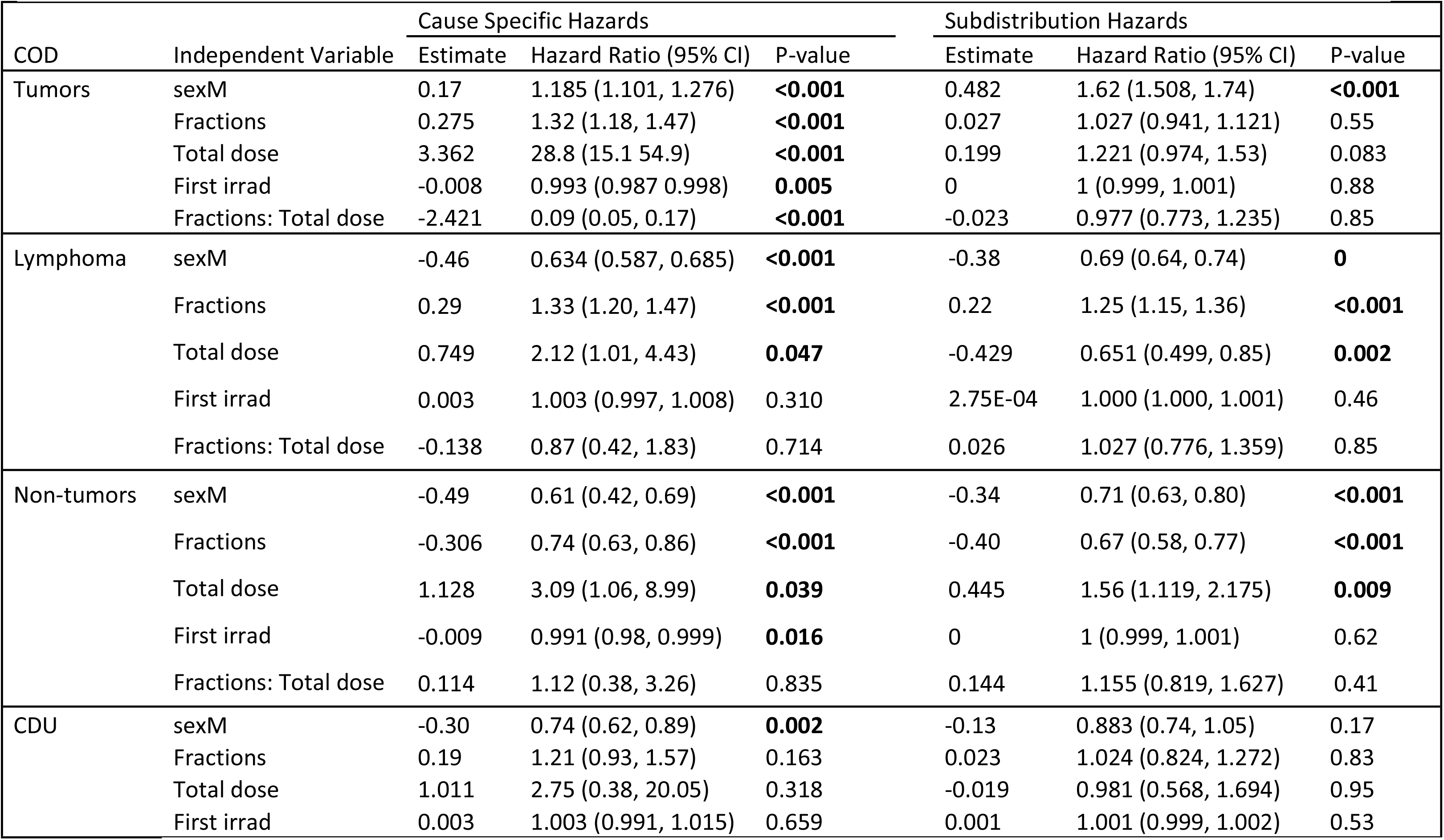
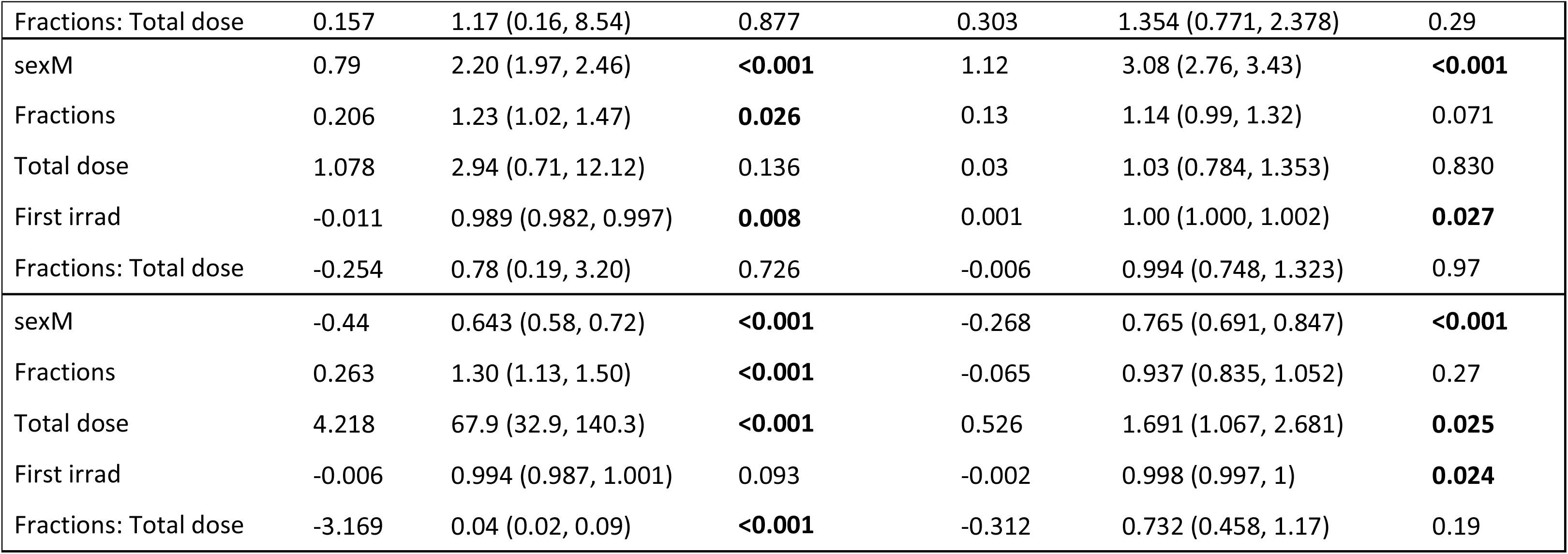
Competing risks model output for cause specific hazards and subdistribution hazards

| COD | Independent Variable | Cause Specific Hazards |  |  | Subdistribution Hazards |  |  |
| --- | --- | --- | --- | --- | --- | --- | --- |
|  |  | Estimate | Hazard Ratio (95% CI) | P-value | Estimate | Hazard Ratio (95% CI) | P-value |
| Tumors | sexM | 0.17 | 1.185 (1.101, 1.276) | **<0.001** | 0.482 | 1.62 (1.508, 1.74) | **<0.001** |
|  | Fractions | 0.275 | 1.32 (1.18, 1.47) | **<0.001** | 0.027 | 1.027 (0.941, 1.121) | 0.55 |
|  | Total dose | 3.362 | 28.8 (15.1 54.9) | **<0.001** | 0.199 | 1.221 (0.974, 1.53) | 0.083 |
|  | First irrad | -0.008 | 0.993 (0.987 0.998) | **0.005** | 0 | 1 (0.999, 1.001) | 0.88 |
|  | Fractions: Total dose | -2.421 | 0.09 (0.05, 0.17) | **<0.001** | -0.023 | 0.977 (0.773, 1.235) | 0.85 |
| Lymphoma | sexM | -0.46 | 0.634 (0.587, 0.685) | **<0.001** | -0.38 | 0.69 (0.64, 0.74) | **0** |
|  | Fractions | 0.29 | 1.33 (1.20, 1.47) | **<0.001** | 0.22 | 1.25 (1.15, 1.36) | **<0.001** |
|  | Total dose | 0.749 | 2.12 (1.01, 4.43) | **0.047** | -0.429 | 0.651 (0.499, 0.85) | **0.002** |
|  | First irrad | 0.003 | 1.003 (0.997, 1.008) | 0.310 | 2.75E-04 | 1.000 (1.000, 1.001) | 0.46 |
|  | Fractions: Total dose | -0.138 | 0.87 (0.42, 1.83) | 0.714 | 0.026 | 1.027 (0.776, 1.359) | 0.85 |
| Non-tumors | sexM | -0.49 | 0.61 (0.42, 0.69) | **<0.001** | -0.34 | 0.71 (0.63, 0.80) | **<0.001** |
|  | Fractions | -0.306 | 0.74 (0.63, 0.86) | **<0.001** | -0.40 | 0.67 (0.58, 0.77) | **<0.001** |
|  | Total dose | 1.128 | 3.09 (1.06, 8.99) | **0.039** | 0.445 | 1.56 (1.119, 2.175) | **0.009** |
|  | First irrad | -0.009 | 0.991 (0.98, 0.999) | **0.016** | 0 | 1 (0.999, 1.001) | 0.62 |
|  | Fractions: Total dose | 0.114 | 1.12 (0.38, 3.26) | 0.835 | 0.144 | 1.155 (0.819, 1.627) | 0.41 |
| CDU | sexM | -0.30 | 0.74 (0.62, 0.89) | **0.002** | -0.13 | 0.883 (0.74, 1.05) | 0.17 |
|  | Fractions | 0.19 | 1.21 (0.93, 1.57) | 0.163 | 0.023 | 1.024 (0.824, 1.272) | 0.83 |
|  | Total dose | 1.011 | 2.75 (0.38, 20.05) | 0.318 | -0.019 | 0.981 (0.568, 1.694) | 0.95 |
|  | First irrad | 0.003 | 1.003 (0.991, 1.015) | 0.659 | 0.001 | 1.001 (0.999, 1.002) | 0.53 |

|  |  |  |  |  |  |  |
| --- | --- | --- | --- | --- | --- | --- |
| Fractions: Total dose | 0.157 | 1.17 (0.16, 8.54) | 0.877 | 0.303 | 1.354 (0.771, 2.378) | 0.29 |
| sexM | 0.79 | 2.20 (1.97, 2.46) | **<0.001** | 1.12 | 3.08 (2.76, 3.43) | **<0.001** |
| Fractions | 0.206 | 1.23 (1.02, 1.47) | **0.026** | 0.13 | 1.14 (0.99, 1.32) | 0.071 |
| Total dose | 1.078 | 2.94 (0.71, 12.12) | 0.136 | 0.03 | 1.03 (0.784, 1.353) | 0.830 |
| First irradiation | -0.011 | 0.989 (0.982, 0.997) | **0.008** | 0.001 | 1.00 (1.000, 1.002) | **0.027** |
| Fractions: Total dose | -0.254 | 0.78 (0.19, 3.20) | 0.726 | -0.006 | 0.994 (0.748, 1.323) | 0.97 |
| sexM | -0.44 | 0.643 (0.58, 0.72) | **<0.001** | -0.268 | 0.765 (0.691, 0.847) | **<0.001** |
| Fractions | 0.263 | 1.30 (1.13, 1.50) | **<0.001** | -0.065 | 0.937 (0.835, 1.052) | 0.27 |
| Total dose | 4.218 | 67.9 (32.9, 140.3) | **<0.001** | 0.526 | 1.691 (1.067, 2.681) | **0.025** |
| First irradiation | -0.006 | 0.994 (0.987, 1.001) | 0.093 | -0.002 | 0.998 (0.997, 1) | **0.024** |
| Fractions: Total dose | -3.169 | 0.04 (0.02, 0.09) | **<0.001** | -0.312 | 0.732 (0.458, 1.17) | 0.19 |

### Lymphoma is the most common COD in neutron irradiated mice while gender significantly impacts probabilities for most COD in control and neutron irradiated mice

**Fig 3A** shows the non-parametric cumulative incidence for each main COD. Lymphoma was the most prevalent, followed by lung tumors, all remaining tumors, non-tumors, and CDU. After dividing the data into control **(Fig 3B)** and neutron-irradiated mice **(Fig 3C)**, we observed that the earliest cases of death occurred around 750 days in both groups of animals. These graphs were also subdivided by gender. For control and neutron-irradiated mice, males had a lower incidence of lymphomas, tumors excluding lung tumors, and non-tumors, but females had a much lower incidence of lung tumors. The difference between males and female mice was significant for both controls and neutron irradiated mice for all causes of death, except CDU (**Fig 3 B** **and** **C**).

**Fig 3:**
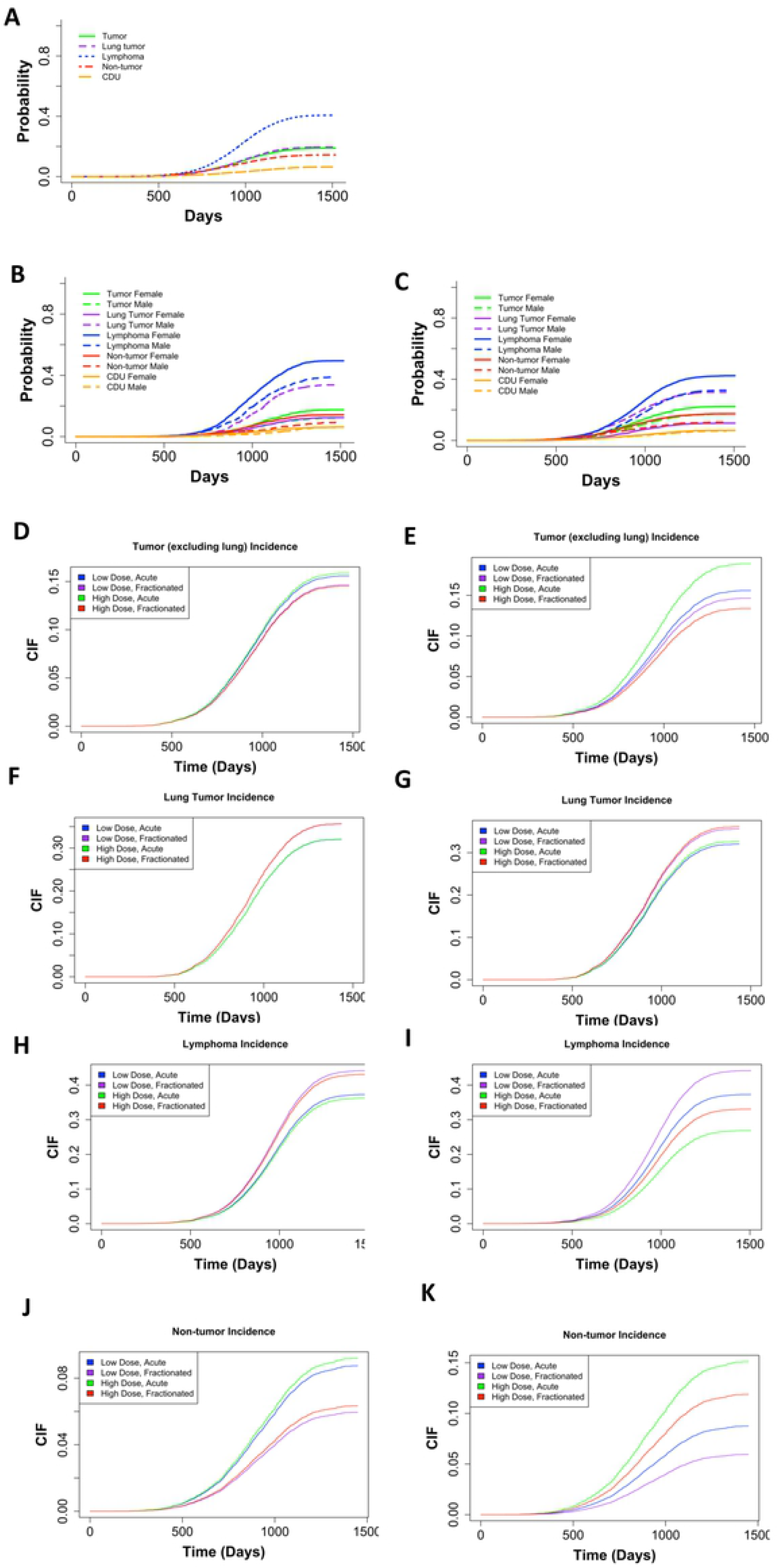
**(A)** Non-parametric CIF for the 5 main categories of COD without grouping. Non-parametric CIF for the 5 main categories of COD grouped my sex for control/sham irradiated mice **(B)** or neutron-irradiated mice **(C)**. P-values for differences in sex for each COD for mice graphed in (B): Tumor = 1.11E-05, lung tumor = 0, lymphoma = 3.75E-11, non-tumor = 2.01E-06, CDU = 0.66, and for mice graphed in (C): tumor = 1.783E-08, lung tumor = 0, lymphoma = 0, non-tumor = 7.07E-12, CDU =.453. Predicted outcome under the following conditions: low dose = 0.01Gy, high dose = 0.1Gy for **(D)** tumors (excluding lung), **(F)** lung tumors, **(H)** lymphomas, and **(J)** non-tumors. Predicted outcome under the following conditions: low dose = 0.01Gy, high dose = 1Gy for **(E)** tumors (excluding lung), **(G)** lung tumors, **(I)** lymphomas, and **(K)** non-tumors. All predicted outputs represent males first irradiated at 120 days. Model output with parameter estimates, hazard ratios (95% confidence interval), and p-value are listed in **Table 2**.

### Tumors other than lung tumors were more frequent with acute neutron exposures, while lung tumors were more common after fractionated exposures

It is important to examine subdistribution hazards, also known as cumulative incidence functions in addition to cause-specific hazards (34, 35, 37–39). The parameter estimates have a less direct interpretation using the Fine and Grey method for CIFs, but explain the overall likelihood of a specific outcome. We included sex, age first irradiated, number of fractions, total dose, and the interaction between fractions and total dose as independent variables in our subdistribution hazards model. The competing risk groups included lymphomas, lung tumors, tumors excluding lung tumors, non-tumors, and CDU. We discovered that females were more susceptible to tumors excluding lung tumors. Specifically, the incidence of tumors excluding lung tumors increased with increase of total dose and decreased with fractionation **(Table 2)**. By graphing predicted outcomes under varying conditions, we found that when there was a tenfold difference in total doses, fractionation was the largest determinant for tumor incidence, with acute exposures resulting in the most tumors **(Fig 3D)**. When the difference in total dose was one-hundredfold, fractionation remained the dominant factor modulating outcome. Acute high dose exposures resulted in the most tumor incidences in all scenarios **(Fig 3E)**.

Examining lung tumors specifically, we found that males were more vulnerable to lung tumors and increases in total dose and fractionation both contributed to increased lung tumor incidence **(Table 2)**. Plotting predicted outcomes under variable conditions, we found that when the difference between high and low total doses was 10-fold, fractionation was the major indicator for lung tumor incidence, with fractionated exposures resulting in the most lung tumors **(Fig 3F)**. When the difference between high and low total doses was 100-fold, fractionation remained the governing influence and fractionated conditions resulted in the greatest lung tumor occurrences **(Fig 3G)**. Under all conditions, high dose fractionated exposures resulted in the highest lung tumor incidences.

### Lymphoma deaths were most frequent in mice that received fractionated low-dose-range neutron exposures

When examining lymphoma deaths, we found that females were at higher risk than males. Increasing the total dose of neutrons decreased lymphoma death incidence, while fractionation increased it **(Table 2)**. Predicted outcome graphs showed that when the difference between high and low total doses was tenfold, fractionation was the most important determinate for increased lymphoma incidence **(Fig 3H)**. Conversely, when the difference between high and low total doses was one-hundredfold, dose became the leading factor and lower total dose exposures resulted in the most lymphoma incidences **(Fig 3I)**. Under all conditions, low dose fractionated exposures resulted in the greatest number of lymphoma incidences.

### Non-tumor deaths were most common after acute high dose exposures

Non-tumor deaths were more prevalent in female mice compared to male mice, increasing with an increase in total dose. Fractionation decreased the probability of death due to non-tumors **(Table 2)**. The predicted outcomes showed that when the difference between high and low total doses was 10cGy, fractionation was the greatest determinate for non-tumor incidence, with acute exposures resulting in the most non-tumor cases **(Fig 3J)**. Conversely, when the difference between high and low total doses was 100cGy, dose became the dominant factor and high-dose conditions result in the most non-tumor death incidences **(Fig 3K)**. Under all conditions, high dose acute exposures resulted in the greatest non-tumor incidence.

### A 60cGy cut-off does not impact CIF outcomes in neutron irradiated mice

Unlike in gamma ray irradiated mice [Zander et al., submitted], no shoulder was observed in the lymphoma or non-tumor incidence graphs for neutron irradiation. Removal of mice exposed to neutrons with total doses of 60cGy or greater did not change the CIF output graphs as it did after removal of mice exposed to high doses of gamma-rays **(S5 Fig)**.

### Mice first exposed to neutrons after 500 days of age began dying later and had increased incidences of death due to tumors

Because of the pronounced difference in longevity of aged mice exposed to neutrons, we examined causes of death in aged mice versus younger animals. Control mice **(Fig 4A)** and gamma irradiated mice **(Fig 4 B-D)** all displayed similar trends in COD patterns with lymphomas as the most prevalent, followed by tumors, non-tumors, and CDU. Control mice, aged gamma-irradiated mice, and mice that were first gamma irradiated at a young age began dying around 600 days **(Fig 4 A, C, D)**. However, data graphs for mice exposed to 6 Gy or more of gamma rays showed a shoulder with increased death incidence beginning around 300 days of age associated with lymphomas and non-tumors **(Fig 4B)**. In gamma irradiated aged mice, there were significantly more cases of non-tumors and significantly fewer cases of lymphoma in comparison to mice exposed at a young age **(Fig 4)**. In all neutron-irradiated mice, solid tumors were more frequent than lymphomas and the effect became more evident when only examining neutron irradiated aged mice (**Fig 4 E** **and** **F**). Aged mice died significantly more often with tumors and non-tumors compared to younger neutron irradiated mice, and there were significantly fewer deaths from lymphomas in aged mice **(Fig 4)**. Mice first irradiated with neutrons when older than 500 days did not begin dying until close to 750 days of age, while mice first irradiated around 115 days began dying around 600 days. Thus, aged neutron-irradiated mice began dying later in life than the control mice **(Fig 4F compared to Fig 4A)**.

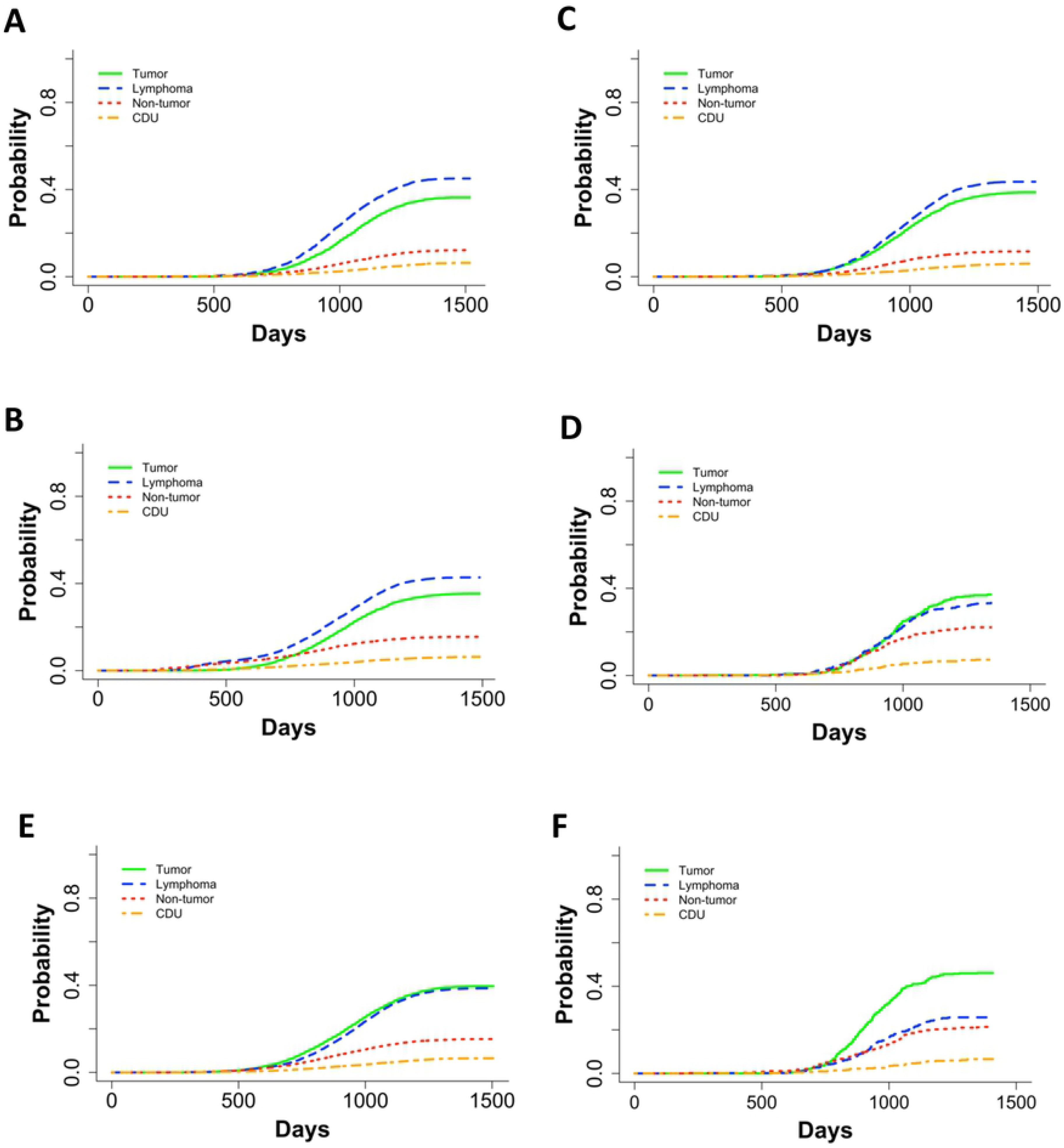

### Gamma and neutron irradiated mice exposed to fractionated once weekly irradiation had a significantly greater death hazard with increase in weekly dose/dose rate

Janus experiments JM8 included mice that were irradiated once a week until death for 45 minutes at a time with dose rates of 0.15, 0.37, or 0.68 cGy/min with cobalt 60 gamma rays or 0.014, 0.035 or 0.056 cGy/min with fission spectrum neutrons. Because the total dose in this experiment was dependent on the lifespan, we performed a separate analysis with Cox PH models using dose rate as our independent variable. We found that as the dose rate increased, the death hazard increased significantly for mice exposed to gamma rays or neutrons **(Fig 5 A and B, and S6 Fig A and B)**. Sensitivity analyses for gamma-and neutron-irradiated mice **(S7 and S8 Fig)** supported these results. We examined KM curves to validate the proportional hazards assumption for using CoxPH models **(S9 Fig)**. The KM curves for sex in neutron-irradiated mice revealed that male and female survival curves overlapped **(S9 Fig D)**. However, when we tested for an interdependence between residuals and time using Schoenfeld residuals **(S9 Fig E)**, we found a non-significant relationship, therefore validating our use of Cox PH models.

**Fig 5:**
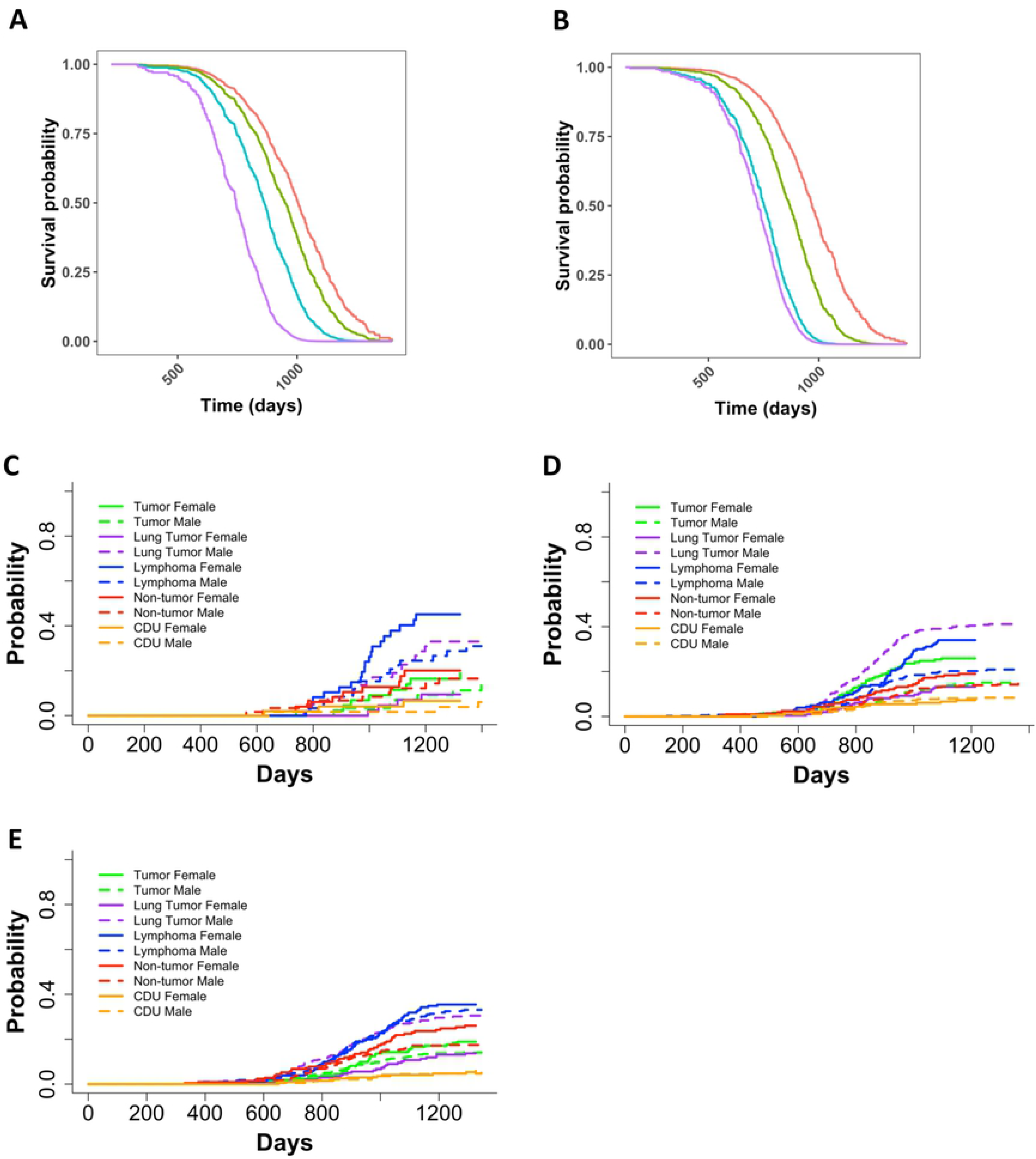
Representative graphs from Cox PH model with age at death as the time scale and sex and categorical dose rate as independent variables for JM8 gamma irradiated mice **(A)** and JM8 neutron-irradiated mice **(B)**. The predicted outcomes shown in A and B represent female mice. Non-parametric CIFs for the five main categories for COD, grouped by sex for JM8 mice that were **(C)** sham irradiated, **(D)** neutron-irradiated, or **(E)** gamma-irradiated. P-values for differences between sexes in **(C)**: tumor = 0.46, lung tumor = 0.0073, lymphoma = 0.085, non-tumor = 0.62, CDU = 0.51, **(D)**: tumor = 7.37E-04, lung tumor = 6.24E-11, lymphoma = 1.04, non-tumor = 0.139, CDU = 0.938 and **(E)**: tumor = 0.18, lung tumor = 5.13E-06, lymphoma = 0.68, non-tumor = 0.029, CDU = 0.57.

### Sex is only significant in once weekly neutron irradiated but not gamma irradiated JM8 mice

In the JM8 experiment, sex played a significant role in neutron-irradiated mice, but not gamma-irradiated mice **(S6 Fig A and B)**. The JM8 dataset consisted of 392 male mice and 215 female mice for neutron exposures and 395 male mice and 215 female mice for gamma exposures. This indicates that sample size is not contributing to a lack of significance for the sex term in our Cox PH model utilizing gamma irradiated mice. Upon further investigation through non-parametric CIFs, we found that control mice only exhibited a significant difference between genders for lung tumors **(Fig 5C).** Neutron irradiated mice had a significant difference between genders for lung tumors, tumors (excluding lung tumors), and lymphomas **(Fig 5D)**. Gamma irradiated mice only displayed a significant difference between lung tumors and non-tumors **(Fig 5E)**. For all treatment groups, females were at a greater risk than male mice for every COD except lung tumors.

## DISCUSSION

Age first irradiated played a significant role in decreasing the death hazard in neutron-irradiated mice. This was evident by examining box plots and the main CoxPH model output for neutron irradiated mice. Age first irradiated was not a significant variable in the main model for gamma irradiated mice **[Zander et al., submitted]**, but it was a significant term in the main model output for neutron-irradiated mice **(Table 1).** A significant interaction between age first irradiated and total dose for neutron irradiated mice was discernible when aged mice data were added to the analysis. As dose increased, the importance of age first irradiated also increased. Among the atomic bomb survivors who were exposed acutely and predominantly to gamma rays, the excess relative risk decreased approximately 21% per decade of age first irradiated (40). In general, it has been accepted that exposures later in life are less risky because the latent phase for slower developing diseases, such as tumors, extends past the anticipated lifespan of someone first exposed to ionizing radiation at an older age. However, the stark contrast in effect for age first irradiated between gamma and neutron irradiated mice indicates that a longer time for disease development is not completely sufficient for increasing risk in younger mice. Because high LET ionizing radiation is more likely to produce clustered DNA damage, which is more difficult to repair (1) and because aged mice are less likely to have efficient cellular recovery mechanisms, it is possible that neutron irradiation late in life may lead to permanent senescence of latent cancer stem cells. Similar hypotheses were stated by others (41, 42) but never in conjunction with neutron exposures. Recently, further investigation into radiation sensitivity and age at exposure have shown that ionizing radiation could make younger populations at greater risk due to disease initiation, but substantially older groups have an increased risk in advancement of preexisting cancerous cells, with an overall bimodal risk distribution with respect to age (43). The large increase in tumor death incidences we observed in neutron irradiated aged mice agrees with this finding. Further work will be necessary to explore these novel hypotheses.

Fractionation decreased the death hazard in neutron irradiated mice **(Fig 1D and Table** Some effects of fractionation on neutron exposures were noted previously (9, 16, 19, 27, 44). A study by De Maio that included a simple 5 fractions fractionation schedule, no significant differences were found for cancer incidence in BC3F1 mice. In a study examining lung cancer risk using the Janus dataset, Heidenreich et al. found that fractionation slightly reduced the relative risk of mice exposed to less than 0.3Gy, but had a detrimental effect at higher total doses (45). We discovered that fractionation was only protective for tumor deaths and did not play a significant role for any other COD group we analyzed **(Fig 2, Table 2, and Supp Fig 4)**. Exploring the mechanism for decreased death hazard through fractionation for mice that died with tumors could lead to potential new radiation treatment or radiation protection strategies.

Lung tumors showed distinct differences in incidence based on gender and response to fractionation. While fractionation was protective against tumor deaths in mice exposed either to neutrons or gamma rays, fractionation was not protective for lung tumors in either case **[Zander, submitted]**. In humans, females exposed to ionizing radiation are at a greater risk for lung tumors compared to males (45, 46). The differences in etiology between lung tumors and all other tumors are worthy of further investigation.

In gamma and neutron irradiated mice, the hazard of death was greater in males compared to females for all causes of death, except for lung tumors. This was one of our most robust findings with cause-specific hazards, subdistribution hazards, and crude CIF results all in agreement for both qualities of radiation. The increased risk for lung tumors in males compared to females, however, may be specific to B6CF1 mice. Reports on lung tumor incidence rates in RFM mice suggested that males exposed to 10 Gy of x-rays were more resistant to lung tumor development than females exposed to 9 Gy of x-rays, the opposite from the trend seen in B6CF1 mice (28, 46, 47). The breathing rate of mice after radiation exposures could potentially contribute to differences in lung tumor susceptibility between genders. It is known that breathing rates increase after high dose radiation exposures, creating a high oxidative state with more free radicals to promote tumor initiation and progression (48, 49). Gender studies examining differences in breathing rates between male and female mice have yet to be conducted, but they could lead to new insights in lung tumor development differences between genders. Tissue samples from mice in the Janus Tissue Archive are available for conducting more detailed molecular biology experiments, such as the PCR based study carried out on Rb and p53 gene deletions in lung adenocarcinomas (50). It is also possible that there is a contribution of genetics to this difference in response since RFM and B6CF1 mice have many genetic differences as one would predict in different mouse strains.

Exploring the differences in disease patterns between aged mice exposed to neutrons or gamma rays led to several significant findings **(Fig 4)**. First, we discovered that lymphoma is the most frequent COD in control mice and gamma irradiated mice, but tumors are the most predominant COD in neutron irradiated mice. This distinction could lead to new hypotheses regarding the differences in disease initiation and development between lymphomas and solid tumors. Second, aged mice exposed to gamma rays died significantly more of non-tumors and died significantly less of lymphomas compared to neutron irradiation survivors. Aged mice exposed to neutrons died significantly more of tumors and non-tumors, and significantly less of lymphomas. The reversal of causes of death for neutrons and gamma ray exposures was surprising. Finally, the non-parametric CIF curves also showed that the age mice began dying was earlier in control and gamma irradiated mice compared to aged neutron irradiated mice **(Fig 4)**. The positive effect for neutrons in tumor deaths appear to be causing major differences in survival, but the mechanisms responsible have yet to be revealed.

Finally, this work included neutron and gamma irradiated mice that were irradiated once weekly until death, making our intention-to-treat method non applicable for these mice (51). Instead, we analyzed these animals using dose rate as the independent variable. Not surprisingly, we found a significant increase in the death hazard as dose rate increased both in gamma and neutron irradiated mice **(Fig 5 A and B)**. A gamma ray exposure of 0.148cGy/min for 45 minutes delivered weekly corresponds to approximately the same total dose one would receive during a computed tomography (CT) scan. Even at this lowest total dose per week, there is a significant decrease in overall survival probability, suggesting that a weekly CT scan would result in harmful health outcomes. Gamma ray exposure resulted in slow decreases in the survival probability at lower dose-rates from 0cGy to 0.048cGy, but changes at higher dose-rates of 0.37cGy/min to 0.68cGy/min had a larger health impact. Conversely, among neutron irradiated mice detriment from weekly irradiation was always significant until it nearly plateaued with little difference between health detriment from 0.035 or 0.056 cGy/min dose rate or weekly neutron doses of 1.58 or 2.52 cGy.

In conclusion, whole body exposures to fission spectrum neutrons evaluated in this work suggest that factors such as age at exposure and period between fractions play a decisive role in the ultimate health outcomes from neutron exposures. Because all of these variations suggest a significant role of the passage of time in resilience to neutron exposure (while physics of neutron exposures occurs instantly on impact) we should turn to the role of biological mechanisms that are played out in whole organisms in our efforts to apprehend the effects of neutron exposures.

## Acknowledgements

The authors of this paper publicly acknowledge the support of Edward Malthouse and Benjamin Haley for their support and statistical experitise. We also thank Carissa Ritner for her support and thoughtful conversations.

## Supplemental Information

**S1 Fig**: Cox PH predicted model output for control mice that received acute or fractionated sham exposures. Independent variables were sex and fractionated status **(A)**, or only fractionated status while stratifying by sex **(C)**. Cox PH model output with parameter estimates, hazard ratios (95% confidence intervals), and p-values for the model in (A) are represented in **(B)** and for the model in (C) are represented in **(D).** The predicted outcome for (A) and (C) represent female mice. **(E)** Kaplan-Meier survival probability for control mice that received that sham exposures as acute vs. fractionated.

**S1 Table**: Description of which mice were removed from our analysis, the corresponding reasoning, and the resulting sample size. N=4130 for control mice and N=10448 for neutron irradiated mice.

**S2 Table:** Causes of death from the Janus dataset categorized by radiation type and subdivided by gender.

**S2 Fig:** Robustness testing for neutron irradiation main model. CoxPH model output from the base model with the following changes: **(A/B)** included mice irradiated at all ages, **(C/D)** included mice irradiated at all ages and an interaction tern between age first irradiated and total dose, **(E/F)** included an interaction term between age first irradiated and total dose, **(G/H)** stratified by sex, **(I/J)** and included exact fraction group and age first irradiated for control samples. The graphs represent predicted outcomes for female mice first irradiated at 120 days.

**S3 Fig:** Kaplan Meier curves showing survival probably vs. time in neutron irradiated and control mice after the filtering shown in Table 1 and removing mice first irradiated after 500 days. KM curves comparing **(A)** sex, **(B)** fractionation status, **(C)** age first irradiated, and **(D)** total dose.

**S4 Fig:** Competing risks models for specific causes of death in neutron irradiated mice with age as a time scale and sex, age first irradiated, total dose, fractions, and the interaction between total dose and fractions as independent variables. Survival curves for cause of death being **(A)** lung tumors, **(B)** all tumors except for lung tumors, and **(C)** non-thymic lymphomas. The graphs represent predicted outcomes for female mice first irradiated at 120 days. **(D)** Model output for non-thymic lymphoma

**S5 Fig:** CIF regression model output using a total dose cutoff of 0.6Gy for tumor (no lung) **(A)**,lung tumor **(C)**, lymphoma **(E)**, and non-tumor deaths **(G)**. Predicted output from CIF model for tumor (no lung) **(B)**, lung tumor **(D)**, lymphoma **(F)**, and non-tumor deaths **(H)**. All predicted outputs represent males first irradiated at 120 days and low dose = 0.01Gy, high dose = 0.1Gy.

**S6 Fig:** Parameter estimates and statistical output from the gamma irradiated model in **Figure 5A (A)** and the neutron irradiated model in **Figure 5B (B)**.

**S7 Fig:** Robustness testing for JM8 gamma irradiation model from figure 5A. CoxPH model output from the base model with the following changes: **(A)** treated dose rate as a continuous variable, **(B)** added age first irradiated, **(C)** treated dose rate as a continuous variable and added age first irradiated, and **(D)** stratified by sex.

**S8 Fig:** Robustness testing for JM8 neutron irradiation model from figure 5B. CoxPH model output from the base model with the following changes: **(A)** treated dose rate as a continuous variable, **(B)** added age first irradiated, **(C)** treated dose rate as a continuous variable and added age first irradiated, and **(D)** stratified by sex.

**S9 Fig:** Kaplan Meier curves showing survival probably vs. time in JM8 mice. KM curves comparing **(A)** dose rate in gamma irradiated mice, **(B)** dose rate in neutron irradiated mice, **(C)** sex in gamma irradiated mice, **(D)** sex in neutron irradiated mice, and **(E)** Schoenfeld residuals from the main model in figure 5B for neutron irradiated mice.

## Notes

Additional Information: Financial support: NIH grants R01OH010469 and RO1CA221150 for all authors.

